# Females use multiple cues to assess competition for egg-laying decisions

**DOI:** 10.1101/2025.01.28.635106

**Authors:** Manvi Sharma, Ashwini Ramesh, Nisha Nagaraj, Kavita Isvaran

## Abstract

Animals deploy a wide suite of behavioural responses to reduce fitness costs related to competition. One common strategy is spatial avoidance of resource patches associated with high competition and this involves a need to accurately assess the level of competition associated with patches. Although well-studied in the context of predation risk (alarm cues vs kairomones), we lack in our understanding of how inputs from multiple types of cues are used for assessing the level of conspecific competition. We tested how *Aedes aegypti* females select sites for oviposition when they encounter multiple types of cues from varying densities of conspecifics. We hypothesised that females perceive conspecific young within these sites as potential competition for their offspring, and tailor their response to the type and level of competition. Through binary choice trials, we assessed female oviposition behaviour in response to a gradient in two types of conspecific cues – egg and larvae. Our results indicate that female response was sensitive to the type of cue. Females showed attraction towards pools with low densities of both egg and larval conspecifics indicating conspecific cueing and the strength of response was stronger towards egg cues. However, this attraction response disappeared at high densities of conspecifics (both egg and larvae) suggesting that the resulting trade-off between benefits from conspecific cueing and costs of competition likely shapes oviposition site selection responses. The broad nature of the oviposition response was similar toward both egg and larval cues, but there were fine level differences in responses towards egg and larvae that depended on the conspecific density level. Our study shows that animals are under selection to use multiple cues to assess conspecific competition to minimise the costs of competition.

## Introduction

Competition from conspecifics for resources can strongly affect lifetime reproductive fitness (Gurevitch et al. 1992; Buxton and Sperry 2017; Ortiz-Jiménez et al. 2021). Animals use a wide suite of behavioural strategies to minimise the negative effects of competition, not only for themselves but also for their kin (Clutton-Brock 1991; Buxton and Sperry 2017). For example, classical studies on Bewick’s swans (*Cygnus columbianus*) show that parents assist their offspring in getting access to food from other families of swans during the winter season when resources are scarce (Scott 1980). In several patch breeding organisms, such as frog and mosquito species, ovipositing females have been shown to avoid pools that have a high density of conspecific offspring, as conspecific competition has been shown to be detrimental to the growth and survival of offspring (Blaustein and Kotler 1993; Schulte and Lötters 2014; Schulte et al. 2020). To deploy such responses effectively, animals need to assess the magnitude of conspecific competition accurately.

In their natural environments, animals are likely to be exposed to multiple cues related to conspecifics that provide information about the magnitude of conspecific competition (Amo et al. 2012; Girard et al. 2015; Deodhar and Isvaran 2018). For example, the density of neighbouring conspecific competitors can be indicative of the amount of unused resources available in the patch (Krause 1992). In addition, information on the quality of the competitors might help assess the level of direct threat (Bro-Jørgensen and Dabelsteen 2008). How individuals use multiple cues has been explored in detail in the context of certain selection pressures, such as predation risk (Ferrari et al. 2008; Luttbeg et al. 2020), mate choice (Rajaraman et al. 2018) or perceived threat of neighbouring ovipositing females (Ramesh et al 2025). For example, prey animals respond to disturbance cues from conspecifics by increasing vigilance but show an escape response towards alarm cues from injured conspecifics (Ferrari et al., 2008). In the context of assessing conspecific competition, we have a limited understanding of how inputs from multiple types of cues are used by animals for decision-making.

We used *Aedes aegypti* as a model to study how individuals seeking a habitat resource respond to multiple types of cues from conspecifics. Manipulating the level of competition by varying conspecific density is logistically challenging and rare (Munga et al. 2006; Belovsky et al. 2011). Instead, to study competition in animals, researchers typically manipulate the magnitude of conspecific competition indirectly, by varying resource quantity (Haché et al. 2013; Svanfeldt et al. 2017). However, animals might respond to cues from conspecific density in a fundamentally different manner when compared to cues from resource quantity, while assessing the level of competition. Our study takes the rarer approach of manipulating densities to understand responses to conspecific competition. Specifically, we examine the selection of oviposition sites by *Ae. aegypti* females in response to two types of cues indicating potential competition levels that their offspring might face: cues related to eggs and larvae present at varying densities.

In the wild, *Aedes* spp. females are likely to face spatiotemporal variation in egg and larval densities when encountering pools for oviposition. For example, early in the mosquito breeding season, females might encounter pools with low densities of larval conspecifics, but egg densities might be very high because of the high density of unhatched eggs from the diapause season (Micieli and Campos 2003). After an event of rainfall, females might encounter a high density of larvae in the pools, but egg densities might be low because most eggs are likely to have hatched following inundation with rainwater. The life cycle characteristics of patch-breeders, such as *Aedes* spp., are peculiar as the larvae are strictly aquatic and incapable of moving from pool to pool. Thus, females are under selection to influence their offspring’s exposure to competition through the selection of oviposition sites and this behaviour is, therefore, expected to be sensitive to the level of conspecific competition among offspring (Resetarits Jr 1996; Kiflawi et al. 2003).

Several challenges have hindered our understanding of the effect of conspecific cues on oviposition decisions. First, there is conflicting evidence on responses of ovipositing females to pools containing conspecific larval density. Most studies suggest that female mosquitoes avoid pools with very high conspecific larval density (Chadee et al. 1990; Edgerly et al. 1998; Kiflawi et al. 2003). High conspecific density in natal patches can reduce fitness in varied ways: by depleting a shared food resource; by interfering with access to a resource; or by releasing growth deterring chemicals. In contrast, a few studies have shown that females are attracted to cues of immature conspecifics while ovipositing perhaps because they use conspecific cues as a proxy for pool quality and thus, copy the choice made by other females (Rudolf and Rödel 2005; Wong et al. 2011; Shragai et al. 2019). Second, there is a lack of studies implementing a rigorous design to test oviposition responses to multiple levels of conspecific densities. Studies have largely examined competition effects and subsequent oviposition site choice response at two levels of conspecific density - low and high and often with indirect cues from competitors (Ower and Juliano 2019; Giatropoulos et al. 2022). Thus, there is a failure in capturing the breadth of female response in a system where there is large variability in oviposition behaviour. Finally, although females encounter spatial and temporal variation in both larval and egg cues in the wild, few studies investigate females’ response to both these types of cues to assess conspecific densities. Larvae present in the pool at the time of oviposition might not share the pool with the eggs laid by the female as eggs hatch only after an episode of rainfall. This is not true for eggs; thus, eggs and larval cues could represent different pieces of information related to competition. As a result, females might be under selection to use cues from both larvae and eggs to assess the conspecific competition and might respond differently to varying egg and larval densities.

Based on previous work and *Aedes* spp. biology and ecology, we hypothesised that ovipositing females should treat conspecifics as representing a competition threat; and tailor their response to the type and level of competitive threat. In the wild, eggs laid by an ovipositing female in a pool hatch together with previously laid eggs as a cohort after inundation by rainfall. Thus, cues from eggs at an oviposition site might indicate the future level of competition that a female’s offspring may face. It is less certain whether larvae present in the pool at the time of oviposition are present when the female’s batch of eggs hatch (upon inundation) because this would depend on when the next rainfall event occurs (Clements 1999; Balasubramanian and Nikhil 2013). Thus, current larvae might not represent the future level of competition as well as do eggs. This peculiar life history characteristics of *Aedes* spp. makes it an interesting model system to study oviposition responses to cues from eggs and larvae. Specifically, we predicted that 1) both larvae and eggs should negatively impact mosquito oviposition at high levels of competition; 2) females should show a dose-sensitive response along the egg density gradient, but not along the larval density gradient because eggs, unlike larvae, represent the future level of competition.

To study the effect of conspecific density on *Ae. aegypti* female oviposition behaviour, we tested female responses to 4 levels of egg density and 4 levels of conspecific larval density in two different experimental setups. To measure the oviposition response of females we used two indices. The first measure, oviposition activity index (OAI), represents the degree to which females discriminate between the treatment and control pools in a binary choice trial and we tested if female preference varied with cue type and density. We also measured total eggs by a female as a measure of pool preference because studies have shown that *Ae. aegypti* lay more eggs in favourable sites and fewer eggs in unsuitable sites by withholding eggs for many days (Trexler et al. 1998; Colton et al. 2003). We find partial support for our hypothesis – females show a dose-sensitive oviposition response to both egg and larval densities. Our study shows that females use multiple types of cues for decision making and that female oviposition responses are complex, both in terms of the amount of investment in reproduction – the number of eggs laid – and the nature of investment – how they distribute eggs between pools.

## Methods

### Study system

We conducted all experiments with individuals from a colony of the *Aedes aegypti* mosquito, maintained in our laboratory at the Indian Institute of Science, Bengaluru. *Ae. aegypti* is a diurnal species, with females typically ovipositing after dawn and before dusk. For discriminating between sites, females have been shown to use short-range gustatory and olfactory cues, such as volatile compounds and chemicals in the pool water (Clements 1992). Typically, the number of eggs laid by a single *Ae. aegypti* from one gonotrophic cycle is below 100 (Woke et al.1956, *unpublished data*). *Ae. aegypti* females are known to deposit eggs, above the water surface, in relatively small ephemeral water bodies, including water-filled containers, tyres, mud pots, rock pools and tree holes (Clements 1999). Competition among immatures and adults in *Aedes* spp. is known to shape many life-history traits, such as body size, longevity, survival and flight potential and studies suggest that *Aedes* females are likely to experience high conspecific densities in the wild (Harrington et al. 2008; Wong et al. 2011).

To test female oviposition response towards varying density of eggs and larvae, we conducted binary choice trials using caged females from the colony. The colony is maintained at a 12:12 hour day and night cycle, at 27 ± 5 °C. The colony was set up in February 2013, with an initial batch of eggs procured from the National Malaria Research Institute. For all our experiments, we used lab-bred females of similar ages as age is known to affect fecundity (Jalil 1974). We conducted our experiments in two steps: 1) Maturation phase and 2) Treatment-assay phase. In the maturation phase, individuals of similar age (7-8 days old) were maintained in cages at a density of 100 females to 50 males. Cages were supplied with *ad libitum* food source, i.e., moist cotton smeared with honey, which was replenished regularly. In the maturation phase, cages were left undisturbed to ensure high-mating frequency in females, followed by artificial blood feeding using a blood-feeding trap (see supplementary information for details on the blood-feeding protocol). Next, in the treatment-assay phase, blood-fed females were isolated using an aspirator and transferred individually to treatment cages where their oviposition behaviour was assayed through binary choice trials.

### Assays for oviposition responses to varying larval density

To measure oviposition responses, a single female was housed in a cage (0.3 m x 0.3m x 0.3 m) and was exposed to two pools – a control pool and a treatment pool – to lay eggs. Based on previous experiments (Sharma et al. 2020), treatment densities of larvae were selected covering the range of densities of larvae likely to occur in pools in the wild. The larval densities selected were – 0, 20, 70 and 155 larvae. Oviposition pools were represented by plastic cups of size 11cm x 4 cm. All cups used were of similar size, shape, colour, and texture because ovipositing mosquitoes are sensitive to container characteristics (Harrington et al. 2008). Treatment cups contained 100ml of freshly prepared water with chemical cues from a particular larval density treatment and control cups contained 100ml of tap water (See supplementary information on cue-water preparation protocol). Both control and treatment cups were lined with strips of filter paper (20 cm x 4 cm) as a substrate for females to lay eggs. The cups were always placed at diagonally opposite ends of the cage and these positions were alternated every trial. The oviposition strips and cue-water/tap water were replenished daily for three days. The three-day period constituted a trial and at the end of each trial, these oviposition strips were dried and eggs were counted to measure female oviposition response. We conducted a total of 112 trials, in 3 blocks, with 35 to 40 trials in each block. Each treatment was represented in every block throughout the experiment.

### Assays for oviposition responses to varying egg density

In the second series of experiments, we measured female response to egg densities. We conducted simultaneous choice trials with blood-fed females using the same protocol that we used for the larval treatment assays. Each cage contained a control cup paired with one egg density treatment (0, 20, 70, and 155). The egg treatment was prepared by attaching a small piece of ovistrip paper (4cm x 2cm) containing the desired number of eggs on the treatment egg strip. The control oviposition cup contained a blank piece of ovistrip paper (4cm x 2cm) attached to the egg strip. Both treatment and control cups contained 100 ml of tap water. Each female was exposed to the assay for three days, and the oviposition strips and tap water were replenished daily. The number of eggs laid by the female on the treatment egg strip was counted by subtracting the treatment egg count from the total egg count. We conducted a total of 60 trials, in 4 blocks with 15 to 20 trials in each block. Each treatment was represented in every block throughout the experiment. <u>Need to mention howOAI was calculated for the control (density = 0).</u>

All experiments were carried out in a laboratory setting using colony mosquitoes between May 2015 – April 2017. We conducted a total of 172 trials. All experiments conducted were in accordance with the Animal Ethics Committee of the Indian Institute of Science.

### Statistical analyses

To understand female decision-making, we tested how female preferences for oviposition sites varied with different densities of egg/larvae by calculating two indices – 1) Oviposition activity index, and 2) Total number of eggs laid by females in both pools.

“Oviposition activity index (OAI)” is commonly used to measure a preference or aversion of breeding habitat in insects (Kramer and Mulla 1979) in binary choice experiments. It is defined as the proportional difference in eggs laid between treatment and control pools. OAI is expressed as (E_t_ – E_c_) / (E_t_ + E_c_) where E_t_ and E_c_ are the numbers of eggs laid in the treatment and control pools respectively, in a given trial. The value of OAI ranges from –1 indicating maximum aversion, to +1 indicating maximum attraction to oviposit in the treatment pool.

In a population of females, females could either distribute their eggs disproportionately between the treatment and control pools or could oviposit all eggs in one of the pools based on the treatment type (egg or larvae). A particular mean OAI could result from females in the population distributing eggs between pools or from the population showing a certain combination of frequencies of females showing a preference and females showing an avoidance for conspecific pools. For example, a mean OAI value of 0 could either indicate that, on average, most females distribute eggs proportionally between the two pools. Alternatively, it could indicate that the population consists of females that have a ‘preference for conspecific pools’, and also females that show ‘avoidance of conspecific pool’, resulting in a mean OAI value of 0. We tested whether the relative frequencies of different oviposition responses – preference, avoidance and distribution – varied when females were faced with eggs vs. larval treatments. To do so, we categorised OAI values greater than 0.8 as a ‘preference response’, OAI values less than −0.8 as an ‘avoidance response’ and values between −0.8 to 0.8 as a ‘distribute eggs response’. We pooled together all trials (regardless of density) for each type of treatment (egg or larvae). We then used Pearson’s chi-squared test (3 by 2 contingency table) with Yates’ continuity correction to test whether the relative frequency of oviposition responses (preference, avoidance, distribution) was correlated with treatment type (egg/larvae). We then formally modelled OAI by fitting a linear mixed-effects model with OAI as the response variable. Type of conspecific cue (egg/larvae treatment) was included as a two-level categorical variable called cue type, density (of egg/larvae) as a four-level categorical variable and the interaction term between cue type and density was included, and the block was included as a random effect. The interaction term was included to test whether oviposition response to conspecific density differed depending on the type of conspecific cue.

In addition, we analysed the total number of eggs laid in both pools as a measure of overall female oviposition response to the treatment. We ran a mixed-effects generalised linear model with Poisson errors to analyse variation in total eggs laid. We included the total eggs laid as the response variable, egg/larvae treatment as a two-level categorical variable called cue type, density as a four-level categorical variable, and the interaction term between density and cue type, and block as a random effect. We also tested if the total number of eggs laid by females in both pools was related to the nature of oviposition response – the strength of avoidance, preference or spreading of eggs between treatment and control pools. We ran a generalised linear model with total eggs laid as the response variable and oviposition activity index (OAI), cue type (egg vs. larvae treatment), and the interaction term between OAI and cue type as the explanatory variables. We also included the quadratic term of OAI and the interaction term of the quadratic variable and cue type in the model. We included the interaction term in the model because we expected female response to vary with the type of treatment (egg vs. larvae) depending on the value of OAI. We used likelihood ratio tests to assess the statistical significance of model parameters. We based our inferences on the full model and dropped only statistically insignificant interaction terms for ease of model interpretation. All analyses were conducted using the statistical software R version 4.0.2 (R Development Core Team 2020).

## Results

Overall, the nature of variation in oviposition response was similar in both egg and larvae assays (Figure 1, Pearson’s χ*2* coefficient = 0.87, df = 2, p-value = 0.65). When we examined variation in female oviposition responses, we found that females showed wide variation in the relative numbers of eggs that they laid in control versus treatment pools, that is, female response varied from avoidance to preference towards conspecific treatments (Figure 1). In the larval assay, 15 % of females laid all eggs in the control pool, 40% of females laid all eggs in the treatment pool and 45% distributed their eggs between the two pools. In the egg assay, 11% of females laid all eggs in the control pool, 35% laid all eggs in the treatment pool and 54% distributed their eggs between control and treatment pools.

**Figure 1:**
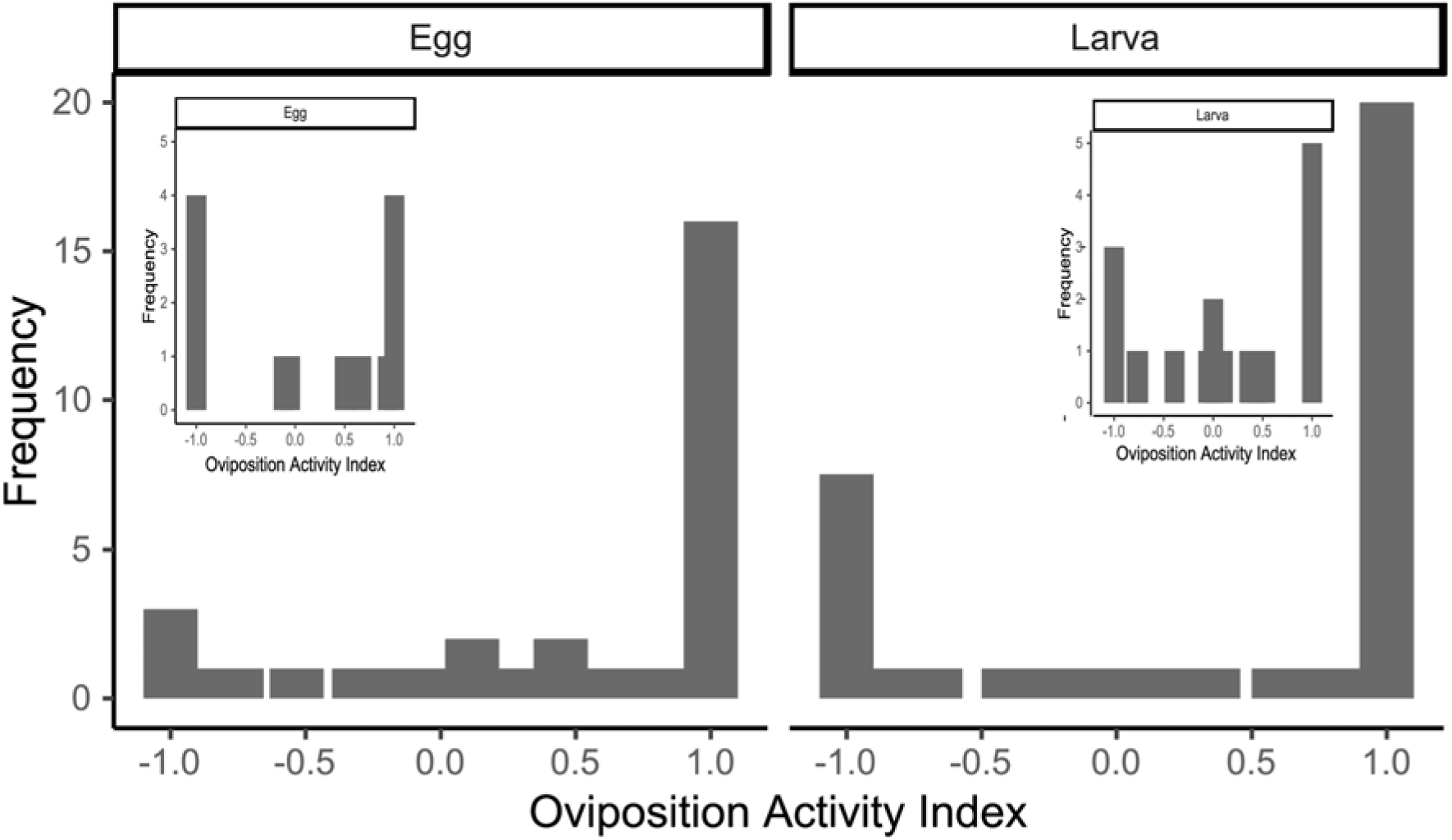
Substantial individual variation in oviposition activity index: In both, egg (left panel) and larval (right panel) assays, most females oviposit all eggs in either control or treatment pool. The figure shows data pooled from three treatments (20, 70, and 155), excluding the control pool (0) treatment. The control pool (0) treatment has been shown in the inset figures for egg and larval assays.

The results from the formal linear mixed-effects model suggest that female oviposition response towards varying levels of conspecific densities was similar towards both types of conspecific cues – larvae and eggs (lmer, cue type term, χ*2*=1.01, df=1, *p*=0.31). For both cues, the oviposition activity index (OAI) was sensitive to conspecific density and decreased with an increase in conspecific density from low to high (lmer, density term: χ*2*=11.93, df=3, *p*=0.007). To interpret differences between density treatments, we used non overlapping 95% bootstrapped confidence intervals (Figure 2).

**Figure 2:**
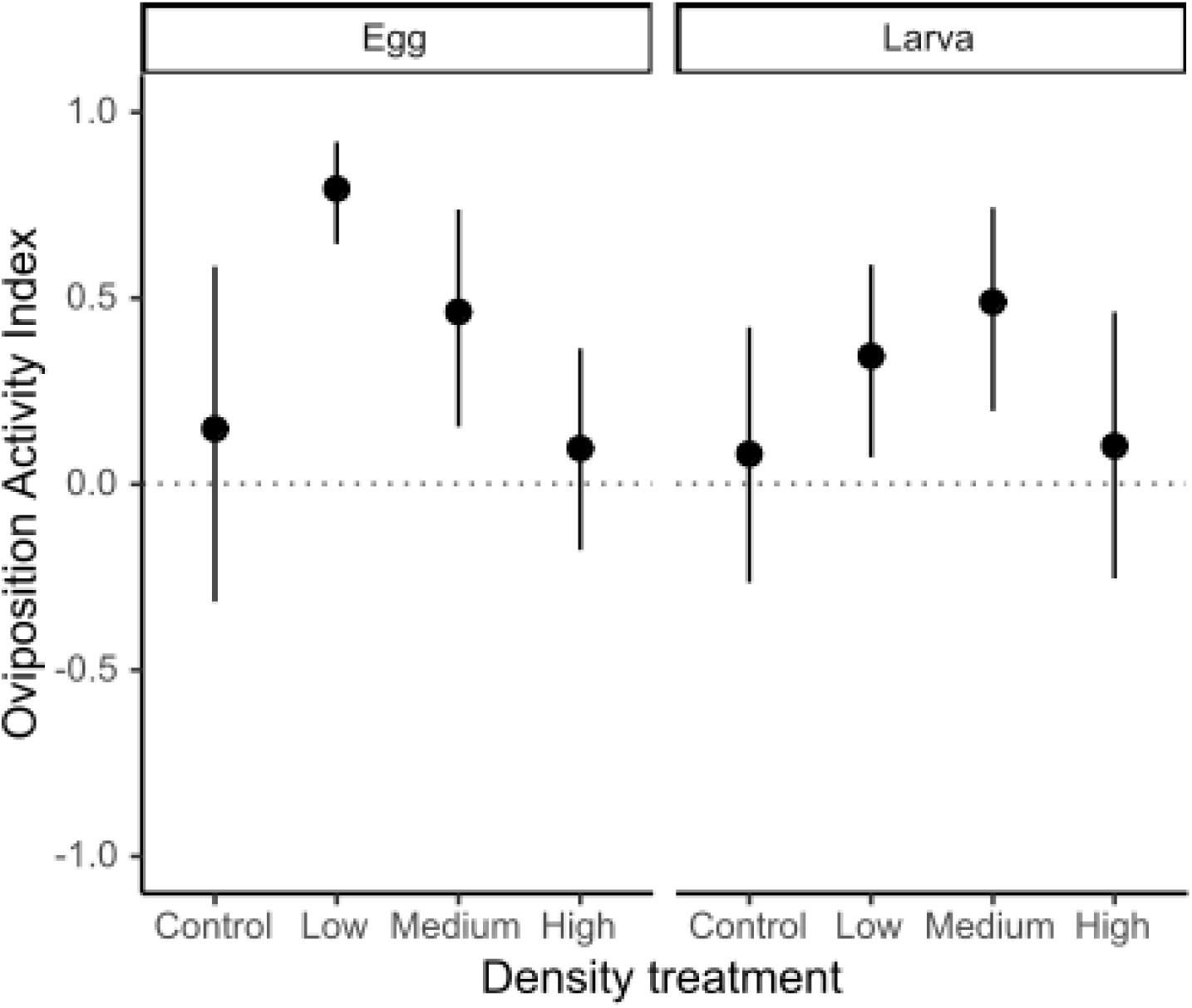
Females show density sensitive oviposition response for both egg and larval cue type. In the egg treatment assay (left panel), females were most attracted to pools with a low density of eggs followed by a medium density of eggs. In the larval treatment assay (right panel), females showed similar levels of attraction towards pools with low and medium larval densities. In both egg and larvae assays, females showed a lack of preference for high densities. Error bars represent 95% bootstrapped confidence intervals on the mean.

For egg density treatments, females showed an attraction response towards both low (20) and medium (70) conspecific densities as indicated by positive OAI values (low density: mean OAI = 0.79, 95% CI = 0.66 – 0.92; and medium density: mean OAI = 0.46, 95% CI = 0.12 – 0.74), and this attraction response was higher for the low density treatment. Similarly, for larval density treatments, females show attraction towards pools with low and medium conspecific densities indicated by positive OAI values (low density: mean OAI = 0.34, 95% CI = 0.09 – 0.59; and medium density: mean OAI = 0.49, 95% CI = 0.19 – 0.74), but this response was similar in magnitude as indicated by the overlap in confidence intervals (Figure 2). The attraction response disappears at high conspecific density of 155 for both egg (mean OAI = 0.13, 95% CI = −0.27 – 0.49) and larval treatments (mean OAI = 0.1, 95% CI = −0.24 – 0.44).

Next, we tested the total number of eggs laid in both pools in response to conspecific densities (Figure 3). Individuals exhibit strong variation in the total number of eggs laid by females per trial, ranging from 168 eggs to 2 eggs. Overall, total eggs laid by females increased with conspecific density, however, this increase was largely in the egg treatment (glmer interaction term, χ*2*=34.21*, df*=3, *p<*0.001, Table 2, Figure 3). We used 95% bootstrapped CI to interpret differences between treatments. In response to egg cues, females laid more than 50% more eggs at the highest conspecific density (155) (mean = 43.2, 95% CI = 27.9 – 59.2) when compared with low conspecific density (20) (mean = 26.7, 95% CI = 18.1 – 36.4) as shown in Figure 3.

**Figure 3:**
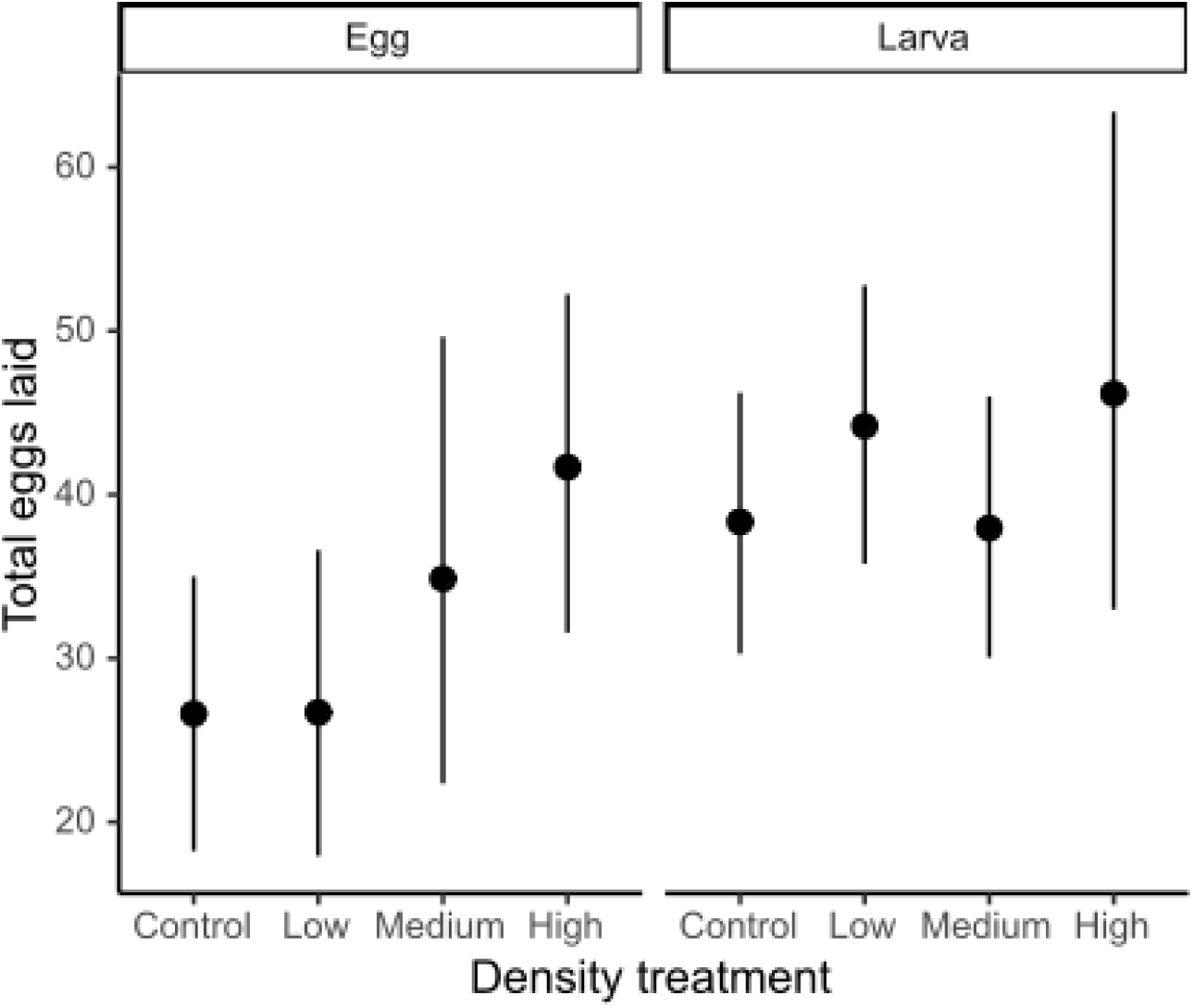
Total eggs laid by a female increased with conspecific density and this increase depended on cue type. Error bars represent 95% bootstrapped confidence intervals on the mean.

**Table 1.** Analysis of variation in the oviposition activity index in relation to conspecific cue (egg/larvae treatment), cue type, density (of egg/larvae), and the interaction term between cue type and density. Likelihood ratio tests are shown for the terms from a linear mixed-effects model fitted to the oviposition activity index.

| Parameter | Coefficient | Std. Error | df | $\chi^2$ | P value |
| --- | --- | --- | --- | --- | --- |
| Intercept | 0.15 | 0.19 |  |  |  |
| (Cue type: Egg, Density: 0) |  |  |  |  |  |
| Density overall effect |  |  |  | 11.93 | 0.007 |
|  |  |  | 3 |  |  |
| Density 20 | 0.64 | 0.27 |  |  |  |
| Density 70 | 0.31 | 0.27 |  |  |  |
| Density 155 | -0.05 | 0.24 |  |  |  |
| Cue type: Larva | -0.07 | 0.25 | 1 | 1.01 | 0.31 |
| Cue type x Density overall effect |  |  | 2 | 2.70 | 0.26 |
| Cue type: Larva x Density: 20 | -0.38 | 0.34 |  |  |  |
| Cue type: Larva x Density: 70 | 0.09 | 0.35 |  |  |  |

**Table 2.** Analysis of deviance in the total eggs laid in relation to cue type, density, and the interaction term between density and cue type. Likelihood ratio tests are shown for the terms from a generalised linear mixed-effects model fitted to the total number of eggs laid.

| Parameter | Coefficient | Std. Error | df | $\chi^2$ | *P* value |
| --- | --- | --- | --- | --- | --- |
| Intercept | 3.37 | 0.08 |  |  |  |
| (Cue type: Egg, Density: 0) |  |  |  |  |  |
| Density 20 | -0.42 | 0.12 |  |  |  |
| Density 70 | 0.18 | 0.35 |  |  |  |
| Density 155 | 0.43 | 0.09 |  |  |  |
| Cue type: Larva | -0.31 | 0.25 |  |  |  |
| Cue type x Density overall effect |  |  | 3 | 34.21 | <0.001 |
| Cue type: Larva x Density: 20 | 0.71 | 0.12 |  |  |  |
| Cue type: Larva x Density: 70 | 0.49 | 0.37 |  |  |  |
| Cue type: Larva x Density: 155 | 0.67 | 0.16 |  |  |  |

Finally, we tested if female oviposition response (avoidance/preference) is correlated with total eggs laid by the female (Figure 4). Females laid fewer eggs in total when they laid all eggs in one pool (control/treatment) but laid a greater number of eggs when they distributed eggs between the two pools (glm OAI squared term, χ*2*=262.8, df=1, *p<*0.001, Figure 4, Table 3). The nature of this response was similar for both egg and larval treatments (glm cue type term, χ*2*=29.71, df=1, *p=*0.1).

**Figure 4.**
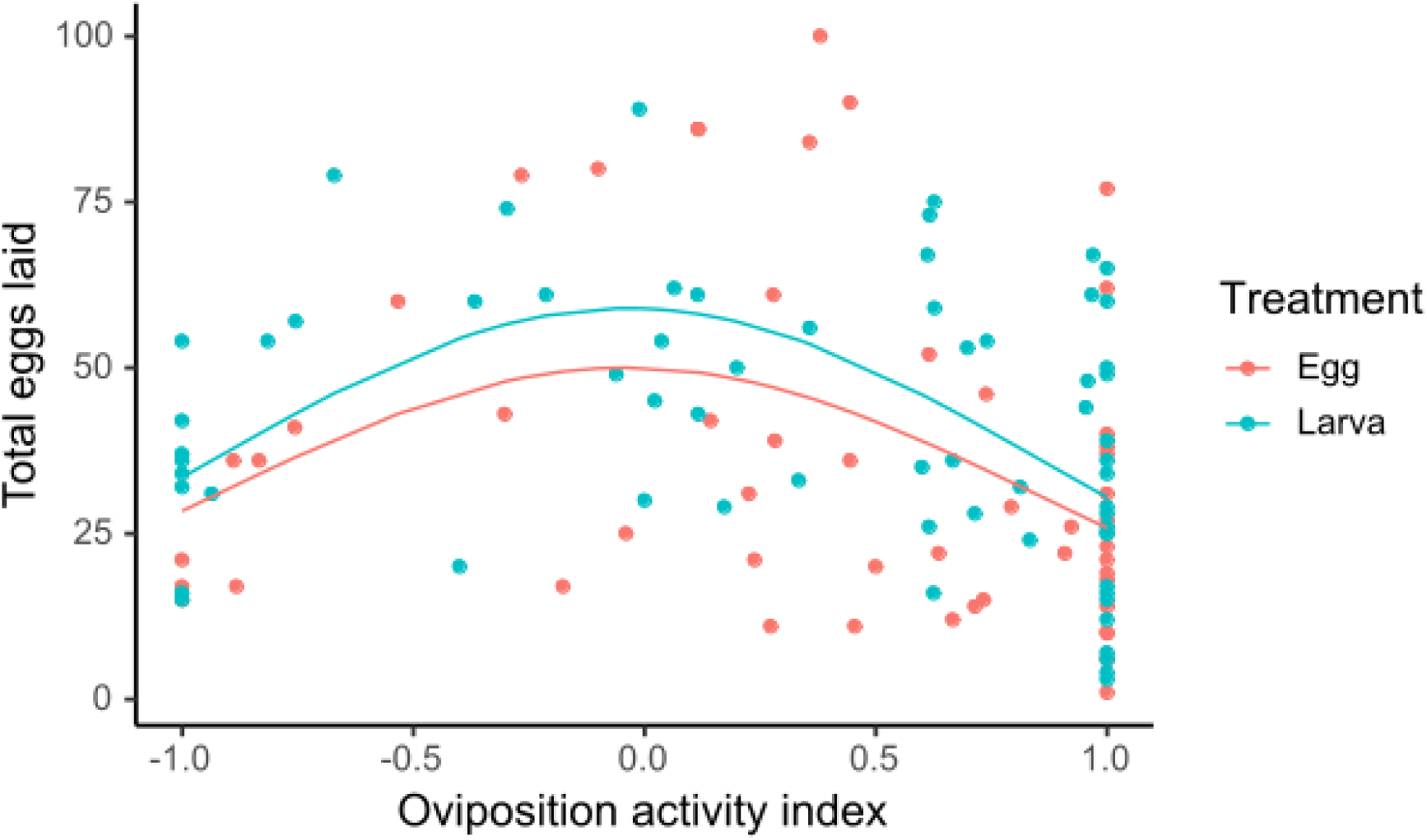
Females laid fewer eggs in total when they laid all eggs in one pool (control/treatment) but laid a greater number of eggs when they distributed eggs between the two pools. Lines represent predictions from the GLMM.

**Table 3.** Analysis of deviance in the total eggs laid in relation to oviposition activity index (OAI), quadratic term of OAI, and cue type. Likelihood ratio tests are shown for the terms from a generalised linear model fitted to the total eggs laid.

| Parameter | Coefficient | Std. Error | df | $\chi^2$ | *p* value |
| --- | --- | --- | --- | --- | --- |
| Intercept | 3.85 | 0.09 |  |  |  |
| (Cue type: Egg) |  |  |  |  |  |
| OAI | -0.06 | 0.07 | 1 | 3.89 | 0.55 |
| OAI<sup>2</sup> | -0.55 | 0.11 | 1 | 262.8 | <0.001 |
| Cue type: Larva | 0.17 | 0.09 | 1 | 29.71 | 0.1 |

## Discussion

How do female responses along a competition gradient compare for different types of conspecific cues? Our study suggest that female response is sensitive to the density of conspecifics – attraction towards low densities and lack of preference at high densities of conspecifics. Females responded towards both types of conspecific young cues – egg and larval cues – suggesting females use multiple cues while assessing pools for oviposition. The broad nature of the oviposition response was similar towards both egg and larval densities, but there were fine level differences in response towards egg and larvae. Females showed a higher level of sensitivity toward egg cues as indicated by the differential response toward low and medium densities. In contrast, females did not discriminate between low and medium conspecific density treatments for larval cues. Overall, our study provides evidence for the hypothesis that females use multiple conspecific cues (egg and larval cues) for oviposition site selection decisions and the level of female sensitivity might depend on the type of cue.

A striking result of our study was the preference that females showed towards the low density of cues from either eggs or larvae. This contrasted with our expectation that females should avoid pools with conspecific cues at all densities. Why are females attracted to low densities of conspecifics? The presence of conspecifics in a pool could indicate a favourable pool without larval predators, or resource-rich pools. By cueing in on conspecifics, females can forego costs associated with sampling pools. This strategy for selecting habitats for breeding has been reported widely from studies on other animals (butterflies: (Raitanen et al. 2014); birds: (Ward and Schlossberg 2004); *reviewed in* Buxton et al 2020). Other mosquito species also exhibit such attraction response (*Aedes albopictus*: (Fonseca et al. 2015); *Culex quinquefasciatus*: (Mokany and Shine 2003)) suggesting that the presence of conspecifics signal the high nutritional quality of habitat. In our study, we compared both larval and egg responses along a gradient of conspecifics and we find that *Ae. aegypti* females are attracted to low densities of both egg and larval conspecifics. This conspecific cueing response disappears when conspecific density increases, perhaps due to the negative effect on larval survival and growth due to high levels of conspecific competition from high-density pools.

Females use multiple cues for decision-making, but how do they respond to these different types of conspecific cues? We hypothesised that eggs represent a reliable estimate of the future risk of competition in the pool. Hence, we predicted that females would show a dose-sensitive response, increasingly avoiding pools along the egg density gradient but not the larval density gradient. Our findings partially support this hypothesis, as females responded to both egg and larvae densities. This suggests that both eggs and larvae might be involved in the assessment of the competition in pools. But there were fine level differences in oviposition response – females preferred low-density pool to medium density pool in egg treatment but not in larval treatment. While both eggs and larvae may indicate habitat stability at low densities, there is a weaker preference for larval treatments in comparison to egg treatments. One explanation for this difference could be mismatch in stage-structure competition. In larval treatments, newly laid eggs would directly compete with late-instar larvae upon hatching, which might outcompete them via starvation for resources. The egg treatments, on the other hand, would result in competition among larvae of the same stage. Future experiments measuring the consequences of this asymmetric competition could improve predictions of female oviposition choices. Another possibility is that females assess competition (in the pool) through egg densities within a different range than they do through larval densities. Currently, few quantitative measurements capture the fitness benefits females gain from ovipositing in pools with varying densities of eggs and larvae. Addressing these gaps can improve our predictions about female oviposition decisions.

*Ae. aegypti* females have been shown to withhold eggs when encountering unfavourable pools (Clements, 1999). Unlike previous studies that investigated *Aedes* spp. oviposition behaviour without monitoring single female behaviour (Harrington et al. 2008; Wong et al. 2011; Albeny-Simões et al. 2014), we conducted experiments with individual females and could measure total eggs laid as an index of oviposition preference. We observed considerable variation in the number of eggs laid by females. Interestingly, the total eggs laid did not depend on the cue type. In both egg and larval treatments, females laid a greater number of eggs when they distributed eggs between pools and laid fewer eggs when they oviposited in either the control or treatment pool. Thus, not only do females decide how many eggs to lay, but this decision also depends on whether they choose to distribute eggs between pools, regardless of cue type. Our findings highlight that female oviposition strategy is complex involving multiple levels of decision-making and reliance on various conspecific cues. For example, the mere presence of neighboring ovipositing conspecifics inhibited egg-laying in larval pools (social inhibition), with females withholding eggs likely in anticipation of future competition (Ramesh et al. 2025). Future research could explore how these different conspecific cues interact to shape the complex oviposition decision-making landscape.

Our results suggest that *Ae. aegypti* 1) has a density sensitive oviposition response to both egg and larval cues 2) shows attraction towards pools with low densities of conspecifics (both egg and larvae) indicating conspecific cueing 3) attraction response disappears at high densities of conspecifics (both egg and larvae) suggesting that competition might be shaping behaviour 4) has a complex oviposition strategy where the number of total eggs laid by females depends on the decision to distribute eggs or put all eggs in a single pool. In the wild, breeding patches are often associated with spatial and temporal heterogeneity in conspecific density and studies show that tracking the presence or performance of conspecifics can be highly beneficial (Andrews et al. 2015; Halliday et al. 2019). In our study, we show that there can be costs associated with co-occurrence in habitats with high conspecific densities. These costs could be in the form of reduced per-capita availability of resources (Mokany and Shine 2003; Stein et al. 2017) or negative effects on growth and reproduction from sharing a habitat with dominant individuals (Bassar et al. 2016). Future work can investigate how females assess the trade-off between the benefits from conspecific cueing and costs of competition to optimize breeding site selection.

## ETHICAL STANDARDS

### Funding

Department of Biotechnology – Indian Institute of Science (DBT-IISc) partnership program; Department of Science and Technology Fund for Improvement of Science and Technology (DST-FIST); Council of Scientific and Industrial Research (CSIR) Grant-in-Aid Scheme no. 37(1636)/14/EMR-II

### Conflict of Interest

The authors declare that they have no conflict of interest.

### Ethical approval

The work we report conforms to the requirements of the Institute Animal Ethics Committee of the Indian Institute of Science, Bengaluru, where the work was carried out, and to the laws of India. This article does not contain any studies with human participants performed by any of the authors.

### Informed consent

For this type of study informed consent is not required.

### Consent for publication

For this type of study consent for publication is not required.

### Statement of authorship

MS (lead), AR, NN and KI conceptualized the study. AR, MS, NN performed the experiment. MS analyzed the data and wrote the first draft of the manuscript. All authors contributed to the final revisions.

## Acknowledgements

We thank Mahalakshmi and Rajashri for maintenance of mosquito colonies; and Karthikeyan Chandrasegaran for helpful discussion on the manuscript.

## Notes

### Competing Interest Statement

The authors have declared no competing interest.

